# Polychrome labeling reveals skeletal triradiate and elongation dynamics and abnormalities in patterning cue-perturbed embryos

**DOI:** 10.1101/2022.12.12.520085

**Authors:** Abigail E. Descoteaux, Daniel T. Zuch, Cynthia A. Bradham

## Abstract

The larval skeleton of the sea urchin *Lytechinus variegatus* is an ideal model system for studying skeletal patterning; however, our understanding of the etiology of skeletal patterning in sea urchin larvae is limited due to the lack of approaches to live-image skeleton formation. Calcium-binding fluorochromes have been used to study the temporal dynamics of bone growth and healing. To date, only calcein green has been used in sea urchin larvae to fluorescently label the larval skeleton. Here, we optimize labeling protocols for four other calcium-binding fluorochromes-alizarin red, xylenol orange, tetracycline, and calcein blue- and demonstrate that these fluorochromes can be used individually or in nested pulse-chase experiments to understand the temporal dynamics of skeletogenesis and patterning. Using this pulse-chase approach we reveal that the initiation of skeletogenesis begins around 15 hours post fertilization, which is earlier than previously thought. We also explore the timing of triradiate formation in embryos treated with a range of patterning perturbagens, and demonstrate that triradiates form late and asynchronously in embryos ventralized via treatment with either nickel at early gastrula stage or with chlorate from fertilization. Finally, we measure the extent of fluorochrome incorporation in triple-labeled embryos to determine the elongation rate of numerous skeletal elements throughout early skeletal patterning and compare this to the rate of skeletal growth in axitinib-treated embryos. We find that skeletal elements elongate much more slowly in axitinib-treated embryos, and that axitinib treatment is sufficient to induce abnormal orientation of the triradiates.

**Highlights:**

- Calcium-binding fluorochromes selectively label the sea urchin larval skeleton
- Pulsed, nested polychrome labeling offers temporal insight into skeletal patterning
- Triradiate formation is delayed and asynchronous in ventralized embryos
- VEGFR inhibition slows skeletal elongation and perturbs triradiate orientation

## Introduction

Skeletogenesis is a highly elaborate process that involves the coordination of multiple cell types to produce and pattern a variety of distinct biomineralized structures within the body plan (Lefebvre & Bhattaram, 2010). However, the molecular mechanisms that underlie skeletal patterning and biomineralization remain poorly understood due to the size and complexity of most vertebrate organisms. The larval skeleton of the sea urchin *Lytechinus variegatus* is an ideal model for studying skeletal patterning due to its relative simplicity. The skeleton is secreted by primary mesenchyme cells (PMCs) that ingress into the blastocoel during early gastrulation and migrate into a ring-and-cords pattern where they fuse to form a syncytium into which the skeletal biomineral is secreted (Hodor and Ettensohn, 1998; Wilt et al., 2008). Calcium is sequestered from the surrounding sea water via endocytosis and is then concentrated in vesicles and deposited as calcium carbonate into the growing spicule (Hu et al., 2020; Schatzberg et al., 2015; Stumpp et al., 2012; Vidavsky et al., 2015; Wilt et al., 2008). This process continues through secondary patterning as the PMCs migrate along the wall of the blastocoel in response to cues from the overlying ectoderm (von Ubisch et al., 1937; Ettensohn and McClay, 1986; Armstrong et al., 1993; Hardin & Armstrong, 1997; Tan et al., 1998; Malinda and Ettensohn, 1994; Ettensohn 1990; Lyons et al., 2012). Our lab and others have identified a number of highly conserved ectodermal cues such as the sulfate transporter SLC26a2/7, sulfated proteoglycans, the TGFβ signal Univin, 5-Lipoxygenase (5-LOX), and vascular endothelial growth factor (VEGF) that are all required for biomineralization and/or skeletal patterning in sea urchin embryos (Adomako-Ankomah and Ettensohn, 2013; Duloquin et al., 2007; Piacentino et al., 2015; Piacentino et al., 2016a; Piacentino et al., 2016b). These cues are also important for patterning and skeletal development in humans and other vertebrates. For example, in humans, SLCa2 is required for cartilage development, skeletal growth, and normal patterning (Hastbacka et al., 1994; Hastbacka et al., 1996a: Hastbacka et al., 1996b; Superti-Furga et al., 1996; Hiala et al., 2001; Kere, 2006). TGFβ signaling plays an essential role in skeletal patterning as well as craniofacial and digit development in mice and humans (Sanford et al., 1997; Wu et al., 2016). Inhibition of 5-LOX, whose metabolites are implicated in human inflammatory disease and in bone resorption, promotes and improves bone regrowth after injury in mice (Haeggstrom & Funk, 2011; Czapski et al., 2016; Chen et al., 2020; Garcia et al., 1996; Traianedes et al., 1998; Cottrell et al., 2013; Biguetti et al., 2020; Simionata et al., 2021). Although human VEGF does not directly impact the skeleton, it is required for the patterning of the limb vasculature, indicating a conserved function in pattern formation (Apte et al. 2019; Eshkar-Oren et al., 2009). Thus, understanding skeletal patterning and biomineralization in sea urchins is likely to provide insight into similar processes in humans and other vertebrates.

An obstacle to understanding the etiology of skeletal patterning in sea urchin larvae is the lack of approaches to readily observe skeleton formation dynamically due to the high motility of the embryos, limiting the appraisal of patterning defects to retrospective examination only. At the end of blastula stage, the sea urchin larva secretes hatching enzymes to degrade the surrounding fertilization envelope (Ishida, 1936; Lepage et al., 1992). Once hatched, the ciliated larva begins to swim and therefore difficult to image without first immobilizing the embryos. To study processes such as gastrulation and PMC migration in sea urchin larvae, researchers have relied on physical barriers such as nylon mesh to reduce movement (Gustafson & Kinnander 1956) or chemical treatments such has 2X artificial sea water to temporarily remove the cilia (Saunders & McClay 2014). However, our attempts to utilize these approaches for live-imaging skeletogenesis have been unsuccessful since the existing methods either do not hold the embryos still enough for confocal imaging of multiple focal planes or do not immobilize the embryos long enough to capture the entire duration of skeletal patterning. Approaches that do adequately immobilize the embryo result in perturbed development and abnormal embryos. Thus, alternative tools are needed to gain temporal insight into skeletogenesis and skeletal patterning.

For decades, calcium-binding fluorochromes such as hematoporphyrin, tetracyclines, calceins, alizarins, and others have been used to label bone and calcifying tissues (Adkins, 1965; Frost et al., 1961; Harris, 1960; Milch et al., 1957; Rahn and Perren, 1970; Rahn et al., 1971). In more recent years, researchers have expanded the use of these chemicals by administering multiple fluorochromes sequentially to investigate temporal dynamics of growth and healing in bones (O’Brien et al., 2002; Pautke et al., 2007). Although some of these fluorochromes have previously been used individually or sequentially to study growth in adult tests of several sea urchin species, calcein green is the only calcium-binding fluorochrome that has been used to fluorescently label the larval sea urchin skeleton (Ellers and Johnson, 2009; Johnson et al., 2013; Kobayashi and Taki, 1969; Rodríguez et al., 2016; Vidavsky et al., 2014). Here, we optimize protocols for larval skeleton labeling with alizarin red, xylenol orange, tetracycline, and calcein blue and establish polychrome labeling techniques with xylenol orange, calcein green, and calcein blue to gain temporal insight into normal and perturbed larval skeletal patterning. We use the approach to reveal that in embryos radialized by NiCl_2_ treatment, triradiate formation is delayed compared to controls and occurs asynchronously. We also show that triradiates are abnormally oriented in embryos in which VEGF signaling is inhibited.

## Results

### Calcium-binding fluorochromes selectively label the sea urchin larval skeleton

We identified five calcium-binding fluorochromes with compatible excitation and emission spectra that permits their simultaneous use: alizarin red (AZ), xylenol orange (XO), tetracycline (TET), calcein green (CG), and calcein blue (CB). Each of these fluorochromes is comprised of at least one fluorescent ring structure as well as one or two calcium-binding moieties (Fig. S1A-E). To optimize these fluorochromes for bright, selective labeling within the sea urchin larval skeleton, we incubated *L. variegatus* embryos in sea water containing each fluorochrome at a range of concentrations from fertilization until the pluteus stage (Fig. S1F-J). At the optimal doses, embryos showed normal skeletal morphology (Fig. 1A) after treatment with fluorochromes, with the exception of alizarin red (Fig. 1B-G). At low doses, the alizarin red signal in the skeleton was faint compared to the level of the punctate background signal. Many embryos incubated with alizarin red exhibited perturbed skeletal patterning (Fig. S1F). When we scored the average prevalence of defects across four biological replicates, we found that alizarin-treated embryos had dramatic skeletal patterning defects even at intermediate doses. This is consistent with previous reports of alizarin toxicity in vertebrate systems (Harris, 1960; Rahn & Perren, 1972; Meyer et al., 2012). The penetrance of these defects was highly variable (Fig. 1B, C), making this mineralization marker unreliable and therefore unsuitable for use in *L. variegatus* larvae. Xylenol orange treatment produced bright and crisp skeletal label at all doses tested, but resulted in minor dorsal-ventral rotational defects at or above 45 µM (Fig. S1G). We therefore selected 30 μM xylenol orange as an optimal dose that selectively labels the skeleton and does not perturb skeletal morphology (Fig. 1B, D; Movie S1). Of all fluorochromes tested, tetracycline produced the faintest signal at all concentrations, even after z-normalization (Fig. S1H), limiting its utility. Calcein green signal was present at all doses, but the skeleton was increasingly difficult to detect amid the non-PMC signal at concentrations of calcein green higher than 30 μM (Fig. S1I). Calcein blue was highly selective for the skeletal spicules, was clearest at 45 μM (Fig. S1J), and did not increase skeletal patterning defects at the optimal dose (Fig. 1B, G). Thus, xylenol orange, calcein green, and calcein blue are each sufficient to selectively label the sea urchin without affecting normal skeletal patterning.

**Figure 1.**
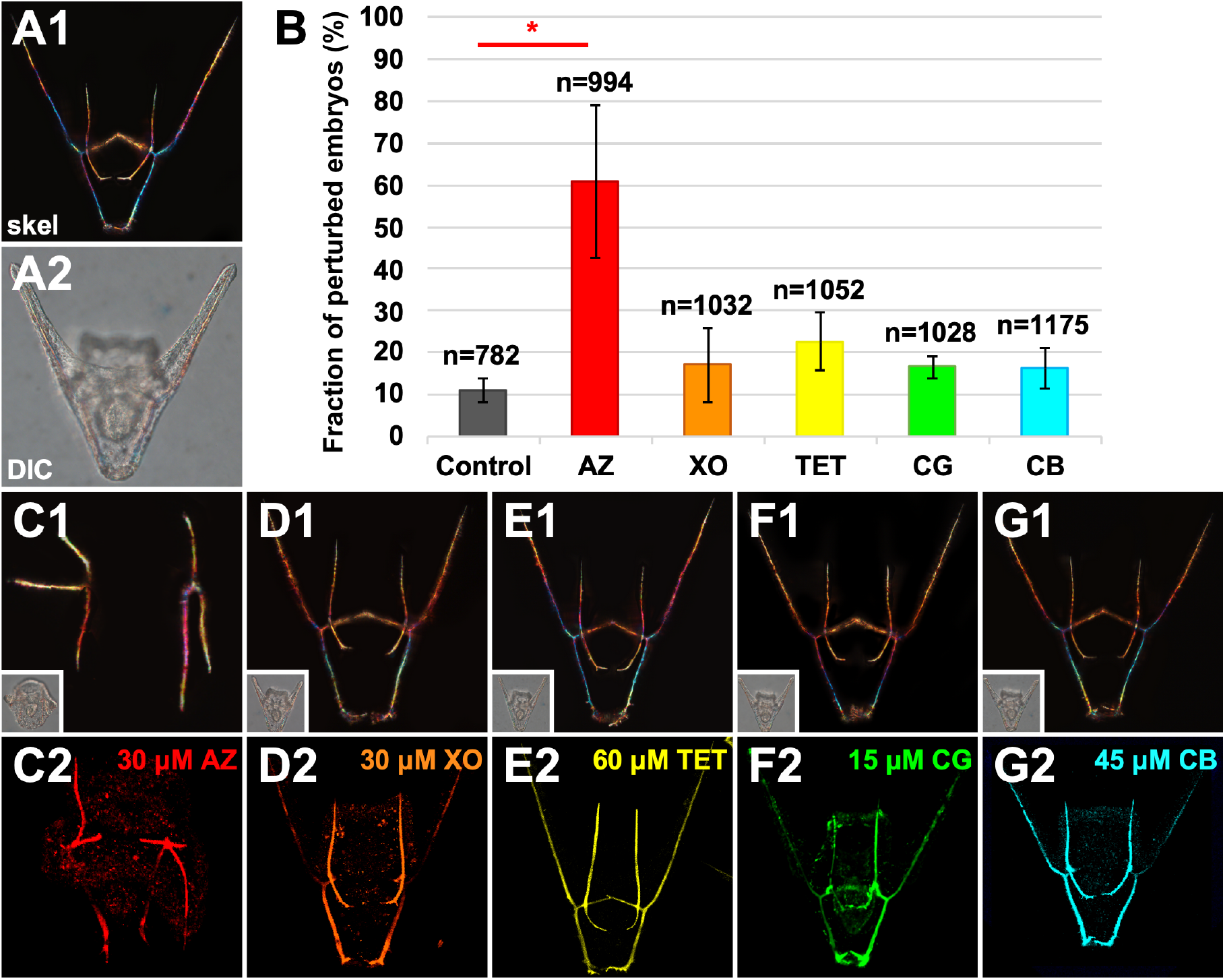
Most of the tested mineralization markers selectively label the larval skeleton without perturbing skeletal patterning. **A**. A control embryo is shown at 48 hpf as skeletal birefringence via illumination with plane-polarized light (1) or as larval morphology via DIC illumination (2) to illustrate normal skeletal pattern and larval morphology. **B**. The fraction of embryos with perturbed skeletal patterning after incubation either in sea water (Control) or in alizarin red (AZ), xylenol orange (XO), tetracycline (TET), calcein green (CG), or calcein blue (CB) is displayed as the average percentage ± S.E.M. from four biological replicates ± SEM; sample sizes (n) are indicated; * p < 0.05 (*t*-test) **C-G**. Skeletal morphology is shown in embryos treated with the indicated fluorochromes as skeletal birefringence (1) and corresponding morphology (DIC, inset), and with the indicated fluorochrome signals (2),

### Polychrome labeling provides temporal insight into sea urchin skeletal patterning

The *L. variegatus* larval skeleton is comprised of two triradiates that appear by 18 hours post fertilization (hpf) that elongate and branch to form the primary and secondary skeletal elements (Fig. 2A) (Lyons et al., 2012; Piacentino et al., 2016a). Historically, our understanding of the temporal dynamics of sea urchin skeletal patterning has been limited by our inability to continuously observe skeletogenesis in real-time within the swimming larvae. Most existing approaches to studying the growth of larval skeletal elements are limited to retrospective or indirect examination that requires fixation of embryos or immunostaining of the PMC syncytium that surrounds the skeletal elements (Ettensohn and Malinda, 1993; Guss and Ettensohn, 1997), and thereby precludes observations and measurement of skeletal dynamics. One exception is a study of skeletal rod elongation dynamics using single pulses of calcein green (Guss and Ettensohn, 1997), which provides some dynamic insights into the elongation of the skeletal elements. Previous studies in other systems were able to infer temporal information about bone growth in live animals using polychrome labeling approaches (Ellers and Johnson, 2009; Nn et al., 1986; O’Brien et al., 2002; Pautke et al., 2007; Saunders et al., 1992; Stuart et al., 1992). To test whether polychrome labeling can be used within the sea urchin larvae to obtain temporal information about skeletal patterning, we sequentially administered two of the selected fluorochromes with the clearest labeling. *L. variegatus* embryos were incubated in xylenol orange for the first 24 hours of development and then incubated in calcein blue for the next 24 hours post-fertilization (hpf) before confocal imaging at 48 hpf (Fig. 2B; Movie S3). The xylenol orange signal was mainly detected in primary skeletal elements, and only faintly labeled the earliest mineralization of the secondary posterior elements (Fig. 2B1). We note that the labeling of the primary VT elements was not easily visible in this exemplar, which is a common outcome in embryos imaged from an anterior viewpoint due to obstruction from the oral hood and/or image depth. The distribution of xylenol orange clearly distinguishes skeletal elements that were formed within the first 24 hours from those that formed later. The calcein blue signal was strongly detected in the secondary skeleton as well as the distal parts of the body rods, which continued to elongate after 24 hours (Fig. 2B2). Calcein blue was also faintly detected in the outermost layers of the primary skeleton (Fig. 2B3-4). This demonstrates that biomineralization of these skeletal elements continues even as other skeletal elements are being formed, resulting in “thickening” of these elements over time, consistent with previous observations (Guss and Ettensohn, 1997).

**Figure 2.**
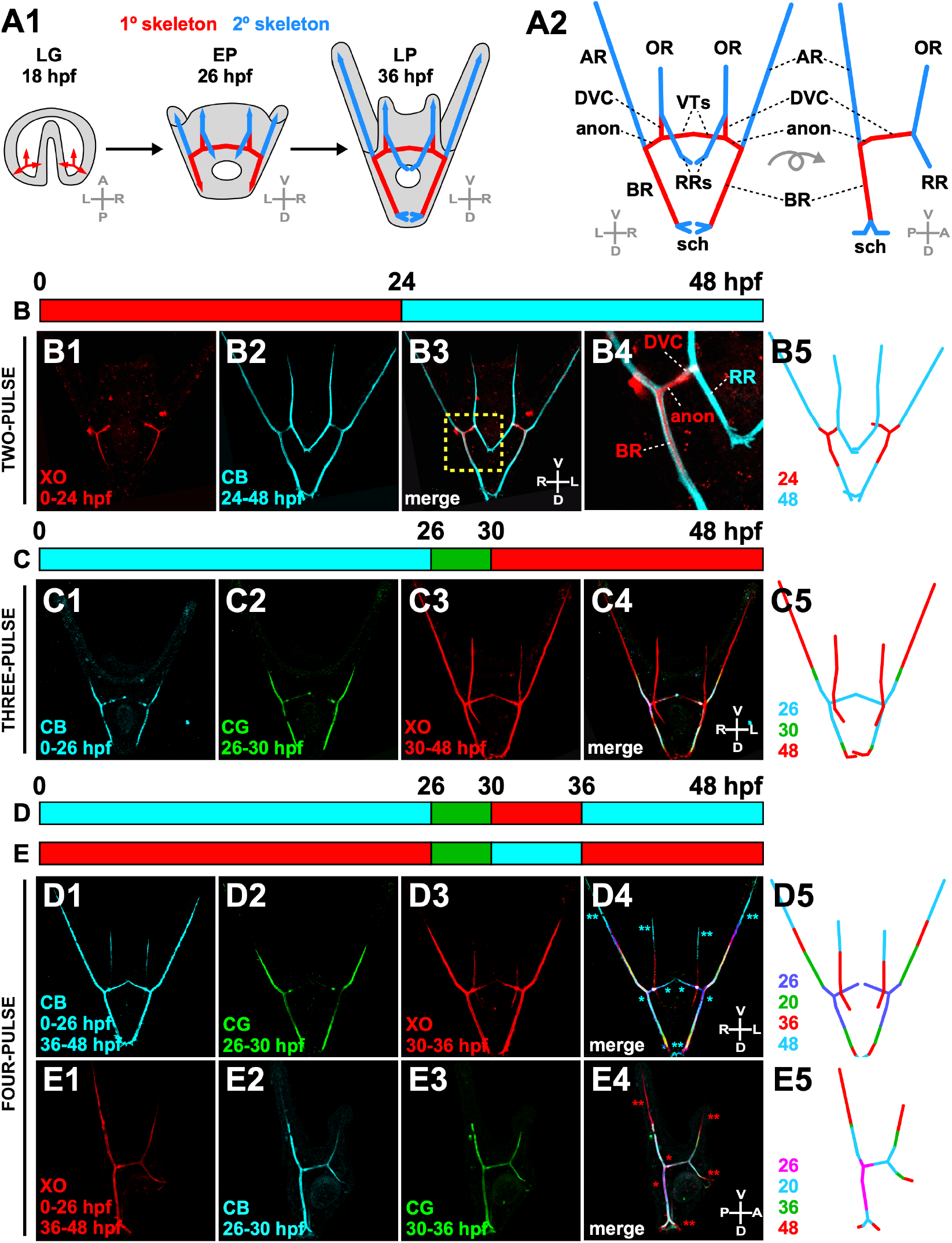
Skeletal growth dynamics are revealed by serial pulse-chase experiments with fluorochrome markers. **A**. The progression of skeletal patterning from late gastrula (LG, 18 hpf) to early pluteus (EP, 26 hpf) to late pluteus (LP, 36 hpf) stages is shown, with primary skeleton (red) and secondary skeleton (blue) labeled (1). Skeletal element identities are indicated in (2); BR body rod; VT ventral transverse rod; DVC dorsal-ventral connecting rod; AR aboral rod; OR oral rod; RR recurrent rod; sch scheitel. **B-E**. The schematics (top) illustrate the experimental protocol, and skeletal schematics illustrate experimental results (5), with new rod extensions indicated. In each case, fluorochromes are shown individually and merged, as indicated. **B**. An exemplar two-pulse embryo is shown at 48 hpf. The indicated section of B3 (yellow box) is enlarged in N4 to show layers of CB label surrounding XO label. **C**. An exemplar three-pulse embryo is shown at 48 hpf. **D-E**. Exemplar four-pulse embryos in *en face* (D) and lateral (E) views are shown at 48 hpf. The first (*) and second (**) non-sequential pulses of calcein blue (D1) or xylenol orange (E1) are spatially distinguishable in merged images (D4-5, E4-5).

The three optimal fluorochromes selected for use in this study have distinct excitation and emission peaks, allowing all three to be used together while still being readily distinguishable (Table S1). To test whether all three could be used sequentially within individual embryos to gain further temporal insight into skeletal patterning without continuous live-imaging, we initially determined the minimum incubation time required for the larval skeleton to have detectable signal when imaged at pluteus stage. To test this, we incubated embryos in one fluorochrome for the first 24 hours of development, then switched to the second fluorochrome for one to four hours before returning to the first fluorochrome for the remainder of secondary patterning (Fig. S2). Calcein blue signal was weakly present in the body rods after as little as one hour of incubation (Fig. S2A1, arrowheads), but was not clearly distinguishable from the background until at least two hours of incubation (Fig. S2A2). Calcein green signal was weakly detectable after one, two, and three hours of incubation, but was brightest and most clearly distinguishable from the background after four hours of incubation (Fig. S2B). Xylenol orange produced detectable signal in the skeleton after one and two hours of incubation, but was brighter and more clearly distinguishable after at least three hours of incubation (Fig. S2C). Thus, to conservatively ensure that we obtained robust and reliable labeling with each fluorochrome, all pulses within subsequent polychrome labeling experiments were performed for at least four hours.

To generate three-color skeletons, embryos were serially incubated in each of the three fluorochromes for at least four hours and then imaged at pluteus stage (Fig. 2C). Since calcein green had the highest level of background signal after longer periods of incubation (Fig. S1I), this fluorochrome was used as the middle, shortest pulse (Fig. 2C2).

The first label, calcein blue, was detected in the primary skeleton as well as the proximal ends of the ARs (Fig. 2C1). The calcein green signal mostly overlapped with the calcein blue signal, suggesting that the majority of the biomineralization that occurred between 26 and 30 hpf served to thicken the existing skeletal elements by increasing their diameter, with only modest elongation of the existing elements evident (Fig. 2C2). Xylenol orange, the final fluorochrome, similarly labeled the thickening of the primary elements as well as the entire secondary skeleton (Fig. 2C3-5).

Skeletal patterning occurs over a 30-hour time period, and it would thus be useful to be able to include more than three pulses of fluorochromes within this timeframe. Since including more than three pulses would require one or more of the fluorochromes to be used repetitively, we next tested the practicality of repeating fluorochromes to increase the number of temporal windows we could query within a single embryo. Four-pulse embryos (Fig. 2D-E) were incubated in the first fluorochrome from 0-26 hpf (Fig. 2D1, E1), the second fluorochrome from 26-30 hpf (Fig. 2D2, E2), the third fluorochrome from 30-36 hpf (Fig. 2D3, E3), and then again in the first fluorochrome from 36-48 hpf. Although the two book-end pulses of the first fluorochrome are not easily discernable within the individual channel (Fig. 2D1, E1), they are spatially distinguishable when all channels are merged based on overlap with other fluorochromes (Fig. 2D4-5, E4-5). Therefore, repetition of fluorochromes is sufficient, in combination with intervening alternate colors, to gain further temporal insight into skeletal rod elongation and thickening.

To gain more refined temporal resolution without shortening the pulse interval below the threshold for detectable fluorochrome signal (Fig. S2), we performed a series of experiments in which we maintained a four-hour pulse and offset the pulse intervals by two hours in different batches of embryos. From these experiments, we could effectively consider two-hour pulses, particularly for length measurements. Using this approach, we obtained a collection of 3-D images from live embryos with fluorochrome switches occurring at regular time points. From these images, we measured the lengths of skeletal elements across multiple time points during skeletogenesis (Fig. 3). We compared our results to those obtained a previous study that obtained lengths from PMC immunostains (Ettensohn and Malinda, 1993) since the resulting PMC membrane label outlines the PMC syncytial cables that enclose the skeletal rods (Piacentino et al., 2016a). However, since these measurements were limited to three timepoints and a subset of the skeletal rods, we also independently determined skeletal rod length measurements via PMC immunofluorescent staining in fixed embryos at a series of timepoints between 20 and 30 hpf. The results show largely comparable results in live and fixed embryos, albeit with some discrepancies, for the majority of skeletal elements (Fig. 3A-H). However, interestingly, the lengths of both the anonymous rods and the DVCs disagreed between measurements in live and fixed embryos (Fig. 3A, C). The anonymous rods and DVCs are each radii of the initial triradiates that terminate in a branch that bounds them, and are relatively static during this interval of skeletal growth. These data suggest that measuring these elements from immunostains overestimates their length, possibly because the precise branch points at each end are ambiguous due to overlapping PMCs within the clusters. The results further suggest that polychrome labeling in live embryos offers a more precise and reliable read-out of the length of the skeletal elements.

**Figure 3.**
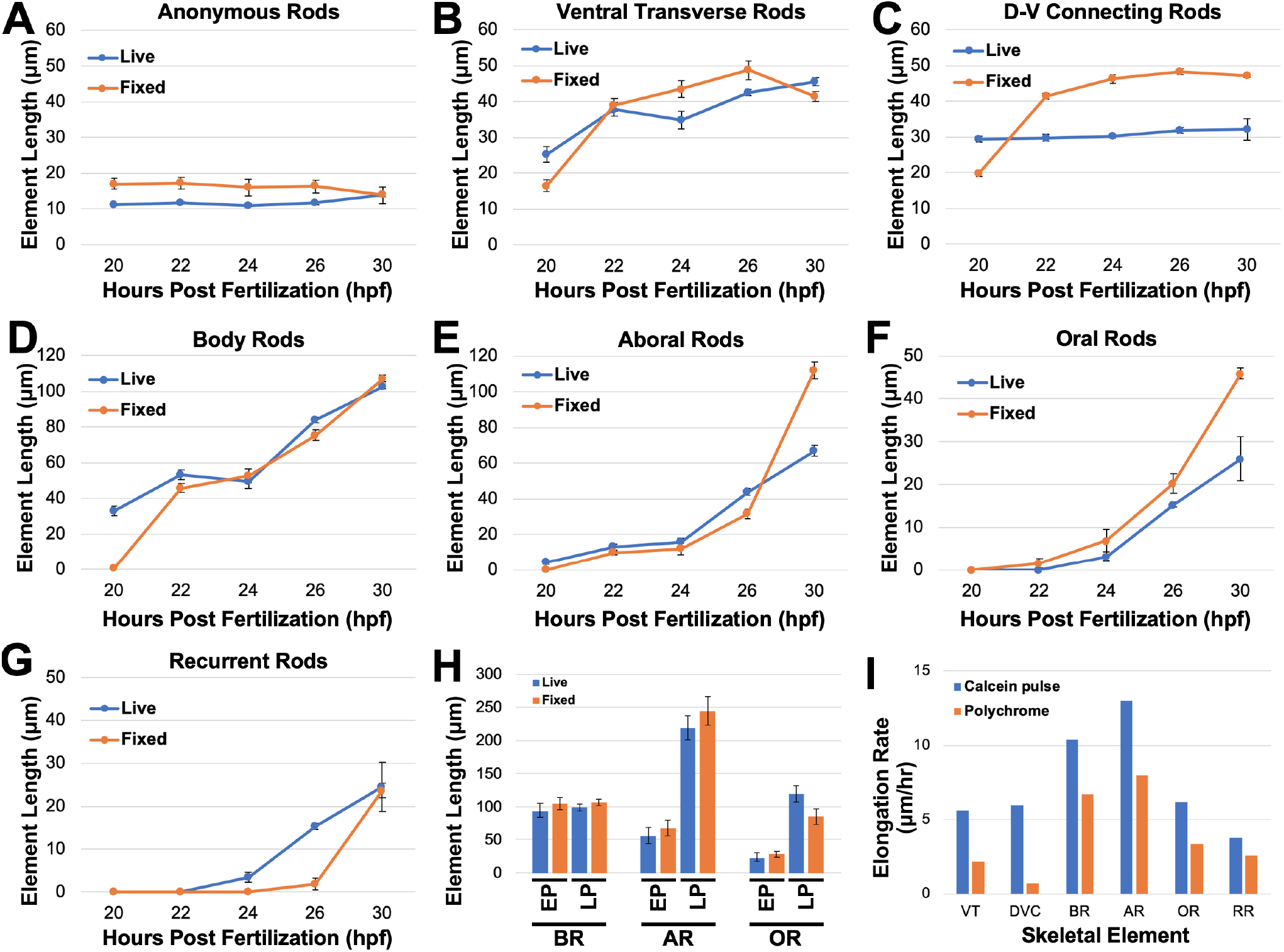
Polychrome labeling is a sensitive approach to studying larval skeletal elongation. **A-G**. The average length of each skeletal rod (in μm) is shown at the indicated time points in live polychrome-labeled embryos (blue) and fixed embryos labeled with a PMC-specific antibody (orange). Error bars show S.E.M. **H**. The average length of the indicated skeletal rods (in μm) is shown at early pluteus (EP, 27-27.5 hpf) and late pluteus (40 hpf) in live polychrome-labeled embryos (blue) and fixed embryos labeled with a PMC-specific antibody (orange, Ettensohn & Malinda, 1993). Error bars show S.D. **I**. Elongation rates (in μm/hr) of the indicated skeletal rods in labeled with a single calcein pulse (blue, Guss & Ettensohn, 1997) or with multiple fluorochromes (orange). Skeletal element names are abbreviated as in Fig. 2.

From these data, we also calculated the average rate of elongation of each skeletal element between 20 and 40 hpf. We compared our results to those obtained in a previous study that measured the elongation dynamics of the different skeletal elements using single pulses of calcein green (Guss and Ettensohn, 1997). Each approach found similar relative elongation rates across the skeletal elements: for example, the body rods and aboral rods elongated most quickly, while the elongation of the DVCs and VTs was slower (Fig. 3I). Interestingly, the elongation rates measured in polychrome-labeled embryos were consistently less than the rates measured in embryos labeled with a single calcein pulse; however, this difference was only about one to five micrometers per hour and could simply be due to technical differences, including 2-D analysis from epifluorescent images using an ocular micrometer in the prior studies (Guss and Ettensohn, 1997), versus 3-D analysis herein from confocal images with precise end-point measurements extracted from the z-stacks. Overall, these results suggest that polychrome labeling captured using confocal microscopy offers the most precise approach for the measurement of skeletal growth dynamics.

### Triradiate formation is synchronous in control and axitinib-, chlorate-, or SB431542-treated embryos but is asynchronous in NiCl_2_-treated embryos

Having established a technique for polychrome labeling of elongating sea urchin larval skeletons, we next explored the temporal information we could gain about skeletal patterning. The earliest event in skeletal patterning is the initiation and elongation of a pair of triradiates. From this initial pattern, primary and secondary skeletal elements branch and elongate (Fig. 2A). The triradiates are thought to be fully extended by the time secondary patterning begins around 24 hpf; so, to assess whether or not triradiate formation proceeded normally in perturbed embryos, we selected 24 hpf as the timepoint at which to perform the fluorochrome switch. Since some of the perturbagens being tested were dissolved in DMSO, we first tested whether DMSO treatment itself produces any disruptions to normal temporal dynamics of triradiate formation. Similar to control embryos whose fluorochromes were switched at the same time point (Fig. 2B), the two triradiates in DMSO-treated embryos appear to be completely labeled with xylenol orange (Fig. 4A, n=16). Taken together, these data suggest that treatment with DMSO alone is not sufficient to perturb the temporal dynamics of triradiate formation.

**Figure 4.**
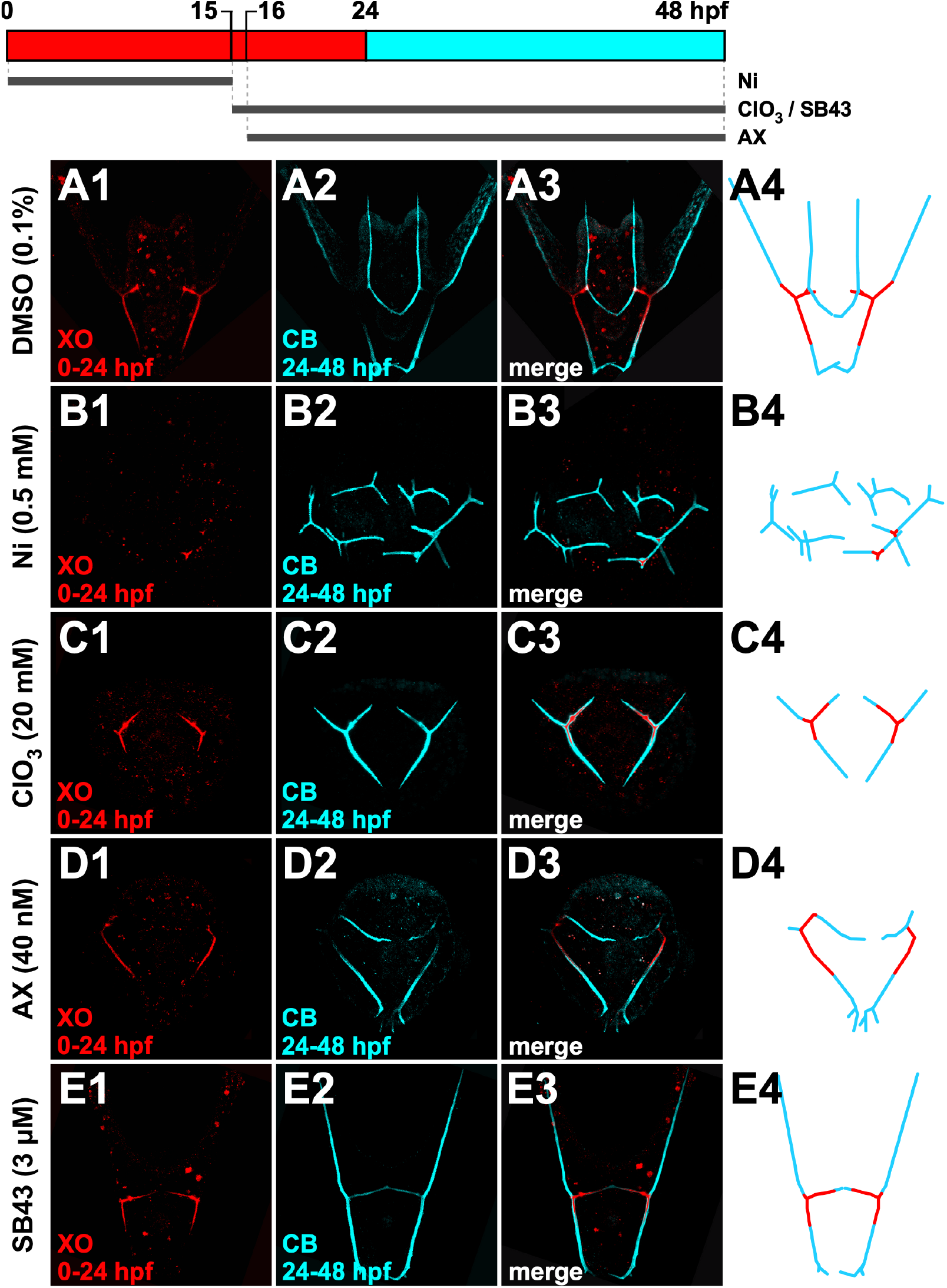
Triradiate formation is asynchronous in Ni-treated embryos but not in ClO_3_-, AX-, or SB43-treated embryos. **A-E**. Embryos that were treated as indicated (A-E) and subjected to two-pulse skeletal labeling (top) are shown at 48 hpf with individual fluorochromes (1, 2) and merged (3) as indicated. Schematics (4) illustrates the results.

Next, we similarly assessed triradiate formation dynamics in embryos treated with four well-established perturbagens that disrupt skeletal patterning: nickel (Ni), chlorate (ClO_3_), axitinib (AX), and SB431542 (SB43). Nickel was one of the earliest chemicals reported to perturb sea urchin skeletal patterning (Lallier, 1956). Since then, researchers have established that nickel treatment disrupts the dorsal-ventral polarity of the ectoderm, resulting in ventralized embryos with radialized skeletons and excess number of triradiates (Armstrong et al., 1993; Duboc et al., 2004; Flowers et al., 2004; Hardin et al., 1992). We were therefore curious to learn whether polychrome labeling would provide any temporal insight into the formation of these extra triradiates. Interestingly, although all nickel-treated embryos had at least 3 triradiates, most embryos (54%, n=46) had produced only one or two triradiates by 24 hpf while the remainder were produced after 24 hpf, as indicated by their labeling only with calcein blue (Fig. 4B). Thus, nickel treatment is sufficient to induce asynchrony in triradiate formation. Furthermore, the xylenol orange label present within the first triradiate(s) appeared to cover a much smaller area than in control embryos. This suggests that less biomineralization had occurred within the triradiates formed by 24 hpf, indicating that nickel treatment is also sufficient to delay in triradiate initiation or elongation compared to controls.

Another perturbagen that affects skeletal development in sea urchin embryos is chlorate, which inhibits the production of sulfated proteoglycans that are normally ventrally enriched (Bergeron et al., 2011; Piacentino et al., 2016a). Embryos exposed to chlorate after gastrulation exhibit skeletal patterning defects, including the loss of ventral elements (Piacentino et al., 2016). When we performed polychrome labeling of embryos that were treated with chlorate at early gastrula stage, we found that both triradiates formed prior to 24 hpf (Fig. 4C). Thus, despite the dramatic stunting of the larval skeleton and loss of many secondary skeletal elements induced by late chlorate treatment, triradiate formation appears to occur synchronously and with normal temporal dynamics in these embryos. The presence of sulfated proteoglycans during gastrulation is therefore not required for timely or synchronous triradiate formation.

Our lab and others have previously investigated the effects of axitinib, a VEGF inhibitor, on skeletal patterning. Treatment with axitinib at early gastrula stage blocks migration of the PMCs out of the ring-and-cords formation, resulting in stunted skeletal rods and loss of some secondary elements (Adomako-Ankomah and Ettensohn, 2013; Piacentino et al., 2016a; Sun and Ettensohn, 2014). We were therefore curious whether axitinib-mediated inhibition of PMC migration might reflect delayed triradiate formation. When we performed polychrome labeling of embryos treated with axitinib at 16 hpf, we found that both triradiates formed by 24 hpf (Fig. 4D; Movie S4). Thus, VEGF signaling after 16 hpf is not required for timely or synchronous triradiate formation.

Lastly, SB43, a specific inhibitor of the TGFβ receptor Alk4/5/7, perturbs anterior skeletal patterning (Piacentino et al., 2015). Although SB43 treatment mainly affects secondary skeletal elements, whether abnormalities in triradiate formation contribute to those phenotypes is an open question. Accordingly, we assessed polychrome-labeled SB43-treated embryos. The distribution of xylenol orange in SB43-treated embryos suggest that triradiate formation is synchronous and occurs prior to 24 hpf (Fig. 4E). Thus, signaling through Alk4/5/7 during or after gastrulation is not required for timely or synchronous triradiate formation.

### NiCl_2_ treatment is sufficient to perturb triradiate timing and synchrony

Of all perturbations explored in this paper thus far, treatment with nickel (Fig. 4B) produced the most interesting perturbations of the temporal dynamics of triradiate formation. To better understand how those dynamics are perturbed, we first performed a more detailed assessment of the normal temporal dynamics of triradiate formation using a series of two-color fluorochrome labeling experiments in which we varied the time of the switch between xylenol orange, delivered first, and calcein blue during early skeletal patterning (Fig. 5A-E). The extent of the xylenol orange signal that appeared in the resulting skeletons corresponds with the extent of triradiate formation and elongation that had occurred by the time of the fluorochrome switch to calcein blue. We found that switching at 12 hpf produced embryos in which no xylenol orange was detected in the larval skeleton itself; however, some signal appeared in calcium-rich vesicles throughout the embryo (Fig 5A, K). Both triradiates were fully labeled with calcein blue, suggesting that they formed after 12 hpf. Embryos with fluorochromes switched at 15 hpf mostly exhibited puncta of xylenol orange signal in the central point of each triradiate along with labeled vesicles (Fig. 5B arrowheads, K), while a minority of this group (35%, n=17) did not incorporate xylenol orange signal in the triradiates. Embryos with fluorochromes switched at 18 hpf had detectable xylenol orange label in all three radii of the triradiates (Fig. 5C, K), while embryos switched at 21 and 24 hpf exhibited complete labeling of triradiates along with partial labeling of other primary skeletal elements such as the body rods (Fig. 5D-E, K). Taken together, this timecourse suggests biomineralization of the triradiates in *L. variegatus* sea urchin larvae is initiated synchronously at or before 15 hpf in most embryos, and that the radii actively elongate between 15 and 21 hpf.

**Figure 5.**
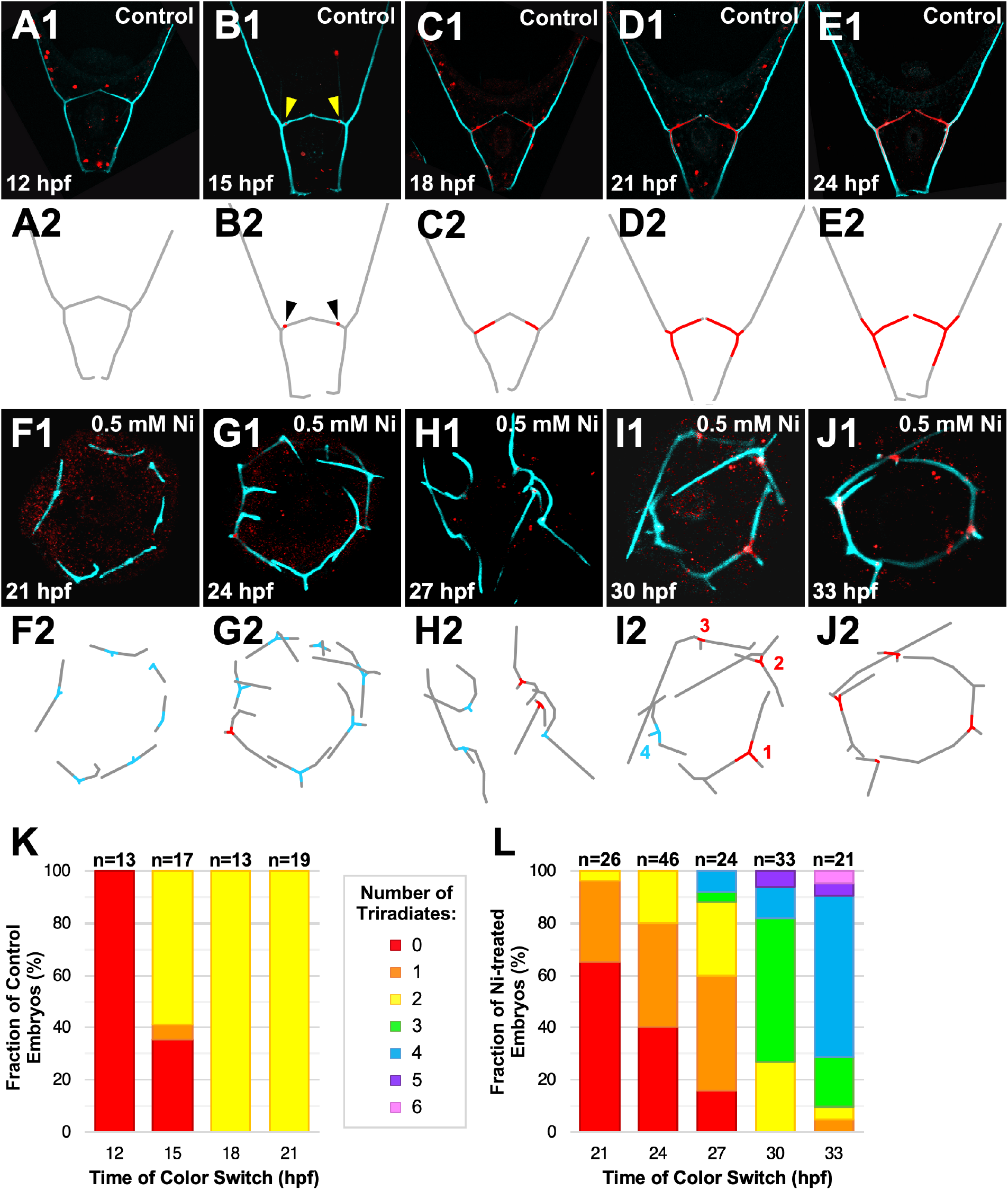
Triradiate formation is delayed and asynchronous in Ni-treated sea urchin embryos. **A-E**. Posterior skeletal exemplars from two-pulse embryos are shown at 48 hpf as merged images (1) and schematically (2). Fluorochromes were switched at 12 (A), 15 (B), 18 (C), 21 (D), or 24 hpf (E). Arrowheads (B1-B2) point to XO label in the center of the triradiates. **F-J**. Ni-treated embryos with fluorochromes switched at the indicated timepoints are shown at 40 to 48 hpf as merged images (1) and schematically (2). Schematics show earliest label detected in each triradiate (red = xylenol orange, cyan = calcein blue) as well as overall skeleton (gray). **J-K**. The average percentage of embryos with the indicated number of triradiates formed at each color switch timepoint is shown for controls (K) and for nickel-treated embryos (L).

We next performed a similar timecourse experiment with nickel-treated embryos in which treated embryos were dual-labeled with xylenol orange first, then with calcein blue, with switches to the second label performed at 21, 24, 27, 30, or 33 hpf. For each color switch time point, we scored the total number of triradiates as well as the number of those triradiates which were labeled prior to the switch, indicating that they had formed before that time point (Fig. 5L). We found that triradiates appeared over a broad range of time points: The first triradiate formed between 21 and 27 hpf (Fig. 5G, L), while the second triradiate formed between 24 and 27 hpf (Fig. 5H, L). The third triradiate formed between 30 and 33 hpf (Fig. 5I, L), while the fourth triradiate formed in nickel-treated embryos at 33 hpf (Fig. 5J, L). In embryos that had five or more triradiates, the fifth triradiates began to appear at 30 hpf and the sixth triradiate at 33 hpf (Fig. 5L). Taken together, these data show that triradiate formation is dramatically delayed in nickel-treated embryos and that triradiate initiation occurs asynchronously over at least a 12-hour interval, compared to controls in which both triradiates are established within a 3-hour window (Fig. 5K-L).

Interestingly, this does not appear to be a nickel-specific effect. Like nickel, chlorate treatment disrupts DV specification and, when treated prior to the onset of gastrulation, produces ventralized embryos with excess numbers of triradiates (Bergeron et al., 2011). When we performed a similar timecourse experiment in which embryos radialized by early chlorate treatment were washed from xylenol orange into calcein blue at 18, 21, 24, 27, and 30 hpf, we found again that the triradiates appeared over a broad range of timepoints (Fig. S3). Like in nickel-treated embryos, subsequent triradiates appeared approximately every three hours (Fig. S3). However, while the first triradiate in chlorate-treated embryos appears later than in controls, its occurrence at 18 hpf is earlier than the first triradiate in nickel treated embryos (Fig. 5G, L; Fig. S3B, F). Thus, ventralization of the ectoderm is sufficient to desynchronize and delay skeletal triradiate formation.

Further analysis of radialized embryos produced another interesting observation: the order in which the triradiates formed in ventralized embryos appeared to be non-random. In the majority (71%, n=21) of embryos that had at least two labeled and two unlabeled triradiates by the time of the fluorochrome switch, the triradiates labeled prior to the time of the fluorochrome switch were adjacent to each other (Fig. 5H-J). Similarly, the majority (81%, n=36) of chlorate-treated embryos with at least two labeled and two unlabeled triradiates by the time of the fluorochrome switch showed that the labeled triradiates had formed adjacent to one another (Fig. S3C). In some cases, the exact order in which the triradiates formed could be inferred based on differences in the extent of xylenol orange incorporation. For example, in Figure 5I the triradiate in the lower right corner of the image has the most expansive xylenol orange label, demonstrating that this triradiate had elongated the most by the time the fluorochromes were switched at 30 hpf. The triradiates then progressively decrease in amount of xylenol orange label counter-clockwise around the image until the final triradiate in the radialized skeleton, which is entirely labeled with calcein blue and therefore formed after the time of the fluorochrome switch. Thus, the initiation of triradiates in ventralized sea urchin embryos appears to be a coordinated process in which subsequent triradiates are initiated near the existing or most recently formed triradiate(s). Future work is needed to explore the mechanism by which this process occurs, and to understand why it becomes desynchronized in ventralized embryos.

### Axitinib treatment is sufficient to significantly reduce elongation rates of skeletal elements and to produce abnormal triradiate orientation

Because the mineralization markers produce fluorescent signal detected using confocal microscopy, the z-slices obtained during image acquisition contain detailed information about the size, shape, and orientation of the skeletal elements. These z-slices can be computationally reconstructed using ImageJ or Napari to produce rotatable 3-D projections of the larval skeleton that are useful for measuring the size and orientation of each skeletal rod (Movie S1; Movie S2). By comparing these features in control and perturbed embryos, we can obtain deeper insight into the etiology and dynamic progression of skeletal patterning defects.

Although we have shown here that axitinib treatment did not perturb the timing or synchrony of triradiate formation (Fig. 4D), previous work by our lab and others has shown that axitinib treatment eliminates or dramatically stunts the elongation of the oral and aboral rods (Adomako-Ankomah and Ettensohn, 2013; Piacentino et al., 2016a). We observed similar defects here (Fig. 4D, compared to Fig. 2B) and were curious to learn whether this stunting was due to differences in elongation rates of the embryos or due to premature termination of skeletal element growth. Thus, we compared the rate of elongation of skeletal elements in control and axitinib-treated embryos during two temporal intervals in which the elements are normally actively elongating (Fig. 3). To test this, we generated three-color control and axitinib-treated embryos that were incubated in xylenol orange from fertilization until 21 hpf, then incubated in calcein green from 21 to 27 hpf, and finally incubated in calcein blue from 27 hpf until imaging at 42 hpf (Fig. 6A, B; Movie S5, Movie S6). We used the resulting rotatable 3-D images to measure the lengths of each skeletal element that were labeled with each of the three fluorochromes to understand the extent to which each element elongated during each incubation period. We found no statistically significant difference between controls and axitinib-treated embryos in the length of most of the skeletal elements at 21 hpf, though we did observe significantly shorter VTs and ARs in axitinib-treated embryos (Fig. 6A-D; Table 1; Table S2). Thus, it appeared that the beginning of skeletogenesis was largely unaffected by axitinib-treatment. However, compared to skeletal elements in control embryos, elements in axitinib-treated embryos elongated much more slowly: by 27 hpf, most skeletal elements in axitinib-treated embryos were significantly shorter than the equivalent elements in control embryos (Fig. 6A-D; Table 1; Table S2). This difference was maintained or increased through 42 hpf, when all skeletal elements except the anonymous rods were significantly shorter in axitinib-treated embryos relative to controls (Fig. 6A-D; Table 1; Table S2). The most dramatic differences in skeletal element elongation were observed in the secondary skeletal elements (Fig. 6C, D). Taken together, these results show that axitinib treatment is sufficient to dramatically slow, but not completely halt, elongation of some primary and all secondary skeletal elements.

**Table 1.**
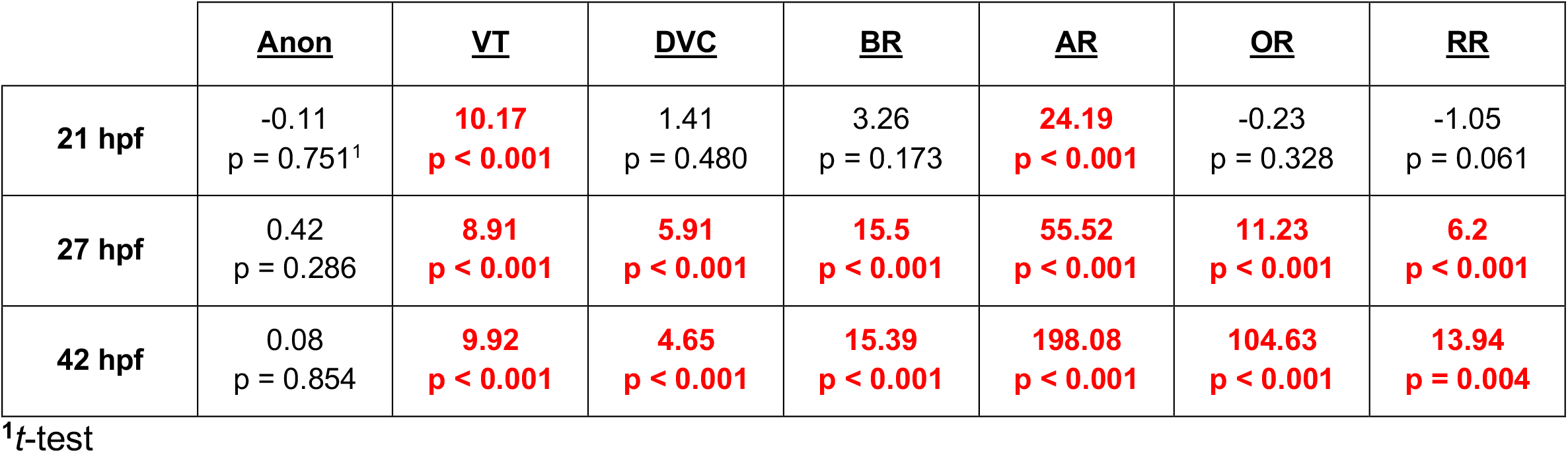
Difference in skeletal element lengths between control and axitinib-treated embryos at three time points.

|  | <u>Anon</u> | <u>VT</u> | <u>DVC</u> | <u>BR</u> | <u>AR</u> | <u>OR</u> | <u>RR</u> |
| --- | --- | --- | --- | --- | --- | --- | --- |
| **21 hpf** | -0.11<br>p = 0.751<sup>1</sup> | **10.17**<br>**p < 0.001** | 1.41<br>p = 0.480 | 3.26<br>p = 0.173 | **24.19**<br>**p < 0.001** | -0.23<br>p = 0.328 | -1.05<br>p = 0.061 |
| **27 hpf** | 0.42<br>p = 0.286 | **8.91**<br>**p < 0.001** | **5.91**<br>**p < 0.001** | **15.5**<br>**p < 0.001** | **55.52**<br>**p < 0.001** | **11.23**<br>**p < 0.001** | **6.2**<br>**p < 0.001** |
| **42 hpf** | 0.08<br>p = 0.854 | **9.92**<br>**p < 0.001** | **4.65**<br>**p < 0.001** | **15.39**<br>**p < 0.001** | **198.08**<br>**p < 0.001** | **104.63**<br>**p < 0.001** | **13.94**<br>**p = 0.004** |

<sup>1</sup>t-test

**Figure 6.**
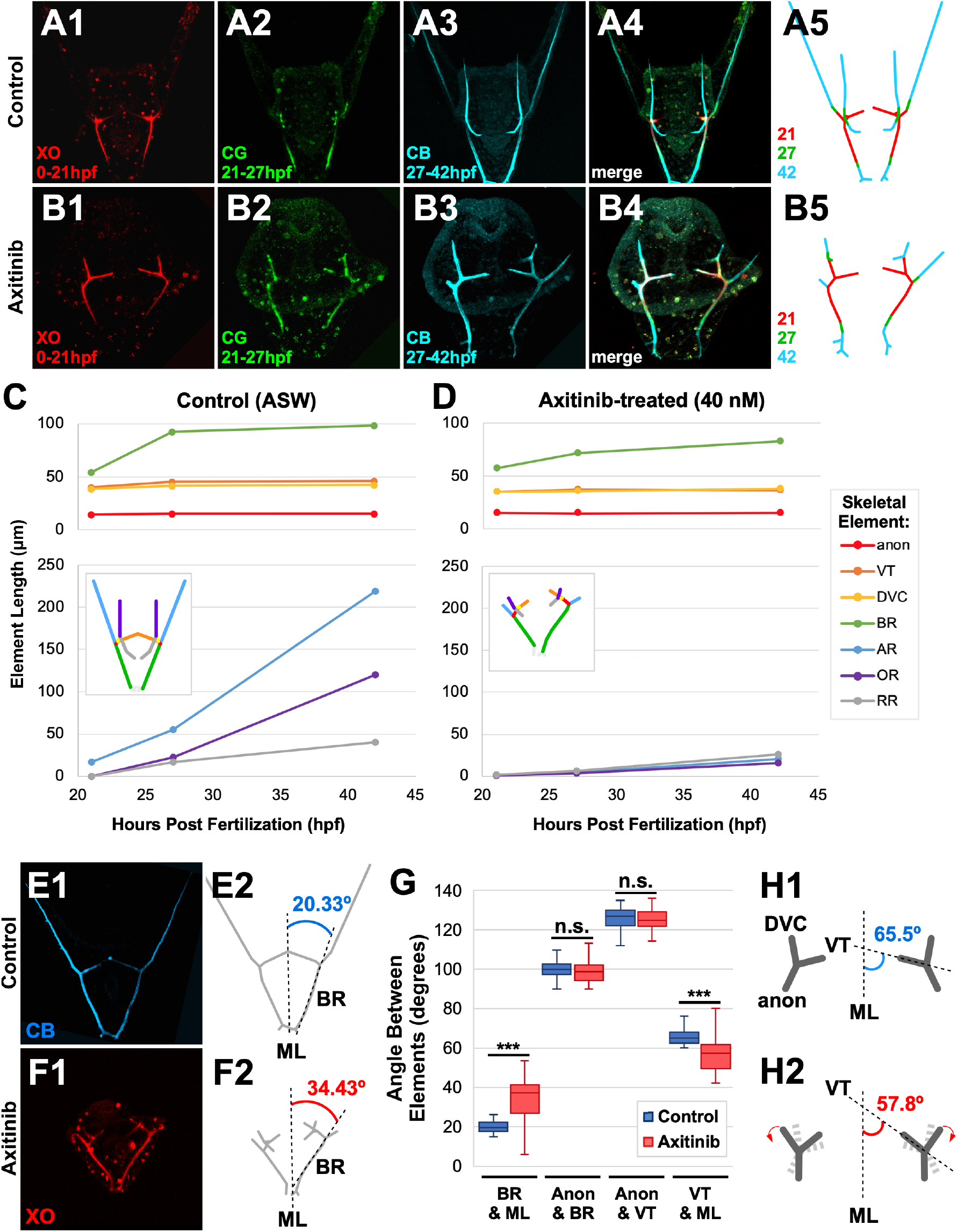
Axitinib treatment perturbs skeletal rod elongation rates and triradiate rotation in sea urchin embryos. **A-B**. A control exemplar (A) and an axitinib-treated exemplar (B) triple-labeled as indicated are shown as individual fluorochromes (1-3), merged (4), and as schematic (5). **C-D**. The average skeletal rod length (μm) per time is shown in control (C, n=15) and axitinib-treated (D, n=12) embryos at 21, 27, and 42 hpf. Skeletal element names are abbreviated as in Fig. 2, and color-coded on inset schematics. **E-F**. The angle between the body rod (BR) and the midline (ML) was measured from posterior views of the labeled skeleton in controls (E) and axitinib-treated embryos (F). **G**. Average angle measurements between the indicated skeletal elements or the midline (ML) are shown as box- and-whiskers plots, with the whiskers reflecting the 10^th^ and 90^th^ percentiles; n ≥ 10; *** p < 0.0001, n.s. not signficant (*t*-test). **H**. Schematics show abnormal triradiate rotation in axitinib-treated embryos (2) compared to controls (1).

Our lab has also previously noted that axitinib-treated embryos have a pronounced anterior-posterior rotational defect, indicated by abnormal orientation of the body rods, in which they are parallel rather than angled (Piacentino et al., 2016a). Here, we used the rotatable 3-D images of fluorochrome-labeled embryos to quantify this difference and found that the angle between the body rods (BR) and the midline (ML) is significantly expanded in axitinib-treated embryos compared to controls (Fig. 6E-H), demonstrating that the body rods in axitinib-treated embryos are dramatically rotated relative to the midline. We were curious to learn whether this rotational defect was a result of structural changes in the shape or branching of triradiates or from overall rotation of the triradiates themselves. Thus, we measured the angles between several pairs of skeletal elements or embryonic features in control and axitinib-treated embryos: the anonymous rods and the ventral transverse rods, the anonymous rods and the body rods, and the ventral transverse rods and the midline (Fig. 6G). We found that the structure of the triradiates themselves was normal in axitinib-treated embryos, since there was no statistically significant difference in the angle between the anonymous rods and the ventral transverse rods in control and axitinib-treated embryos. The angle of the branching of the anonymous rods was also unaffected, since there was no statistically significant difference in the angle between the anonymous rods and the body rods in control and axitinib-treated embryos. Thus, axitinib treatment is not sufficient to perturb branching of the anonymous rods. Since the shape and branching of the triradiates is normal but the body rods are abnormally oriented relative to the midline, this suggests that the orientation of the triradiates relative to one another is perturbed in axitinib-treated embryos. To test this hypothesis, we measured the angle between the ventral transverse rods and the midline and found that this angle was statistically significantly smaller in axitinib-treated embryos compared to controls (Fig. 6G, H). Interestingly, we also observed that axitinib-treated embryos had a statistically significant difference in this angle between the left and right triradiates (not shown), suggesting that bilateral symmetry is perturbed by loss of VEGF signaling. These findings suggest that the anterior-posterior orientation defects observed in axitinib-treated embryos are due to abnormal rotation of the triradiates themselves rather than abnormal shape or branching of the triradiate spicules, which in turn suggests that VEGF signaling is required both to align and to coordinate the orientation of the triradiate about the AP axis to produce normally oriented primary skeletal elements.

## Discussion

In this study, we establish polychrome labeling of the sea urchin larval skeleton as an approach for measuring the temporal dynamics of skeletal rod elongation, thickening, and patterning. Previous studies have used calcein to label the sea urchin skeleton and calcium-bearing vesicles (Guss and Ettensohn, 1997; Killian and Wilt, 2017; Schatzberg et al., 2015; Vidavsky et al., 2014; Vidavsky et al., 2015; Winter et al., 2021). Here, we successfully replicate these results, demonstrating strong calcein labeling in the skeleton as well as in vesicles in cells throughout the embryos (Fig. 1F, S1I). We also optimize labeling the sea urchin larval skeleton with four other calcium-binding fluorochromes that have previously been used to label mineralizing tissues in other systems: alizarin red, xylenol orange, tetracycline, and calcein blue (Harris, 1960; Milch et al., 1957; Rahn and Perren, 1970; Rahn et al., 1971). While xylenol orange and calcein blue selectively label the larval skeleton without perturbing normal skeletal patterning (Fig. 1D, G; Fig. S1G, J; Movie S1), other fluorochromes produced faint labeling (tetracycline) or teratogenic effects (alizarin red) (Fig. 1C, E; Fig. S1F, H). These results are consistent with those observed by others: alizarin red solutions have noted toxicity in vertebrate systems when used to label bone structures (Harris, 1960; Rahn & Perren, 1972; Meyer et al., 2012) and tetracycline has previously been observed to produce faint labeling (Johnson et al., 2013).

Calcium-binding fluorochromes have previously been used to gain temporal information about bone growth and regeneration through sequential addition of multiple fluorochromes (Ellers and Johnson, 2009; Nn et al., 1986; O’Brien et al., 2002; Pautke et al., 2007; Saunders et al., 1992; Stuart et al., 1992). However, to date no polychrome labeling experiments have been performed in the developing sea urchin larvae. Here, we optimize a protocol for polychrome labeling of the sea urchin larval skeleton with up to three fluorochromes via nested pulse-chase experiments. We demonstrate that xylenol orange, calcein, and calcein blue are distinguishable within a single larval skeleton and that clear fluorochrome labeling within the larval skeleton can be achieved after at least two hours of exposure to calcein blue, three hours of exposure to xylenol orange, or four hours of exposure to calcein green (Fig. 2; Fig. S2). This timeframe is in agreement with previous reports regarding calcein incorporation rates into sea urchin larval skeletons (Killian and Wilt, 2017).

We also demonstrate that at least four labeling pulses can be included within a single larval skeleton (Fig. 2-E), which is comparable to polychrome labeling approaches used in other systems (Pautke et al., 2005; Solheim, 1974). Using this polychrome labeling approach, we gain temporal information about when the various elements were formed based on where each fluorochrome is incorporated within the skeleton. This provides distinct retrospective insight into the temporal dynamics of skeletal elongation during multiple time intervals in live embryos. We also compare this polychrome labeling approach to traditional approaches using PMC immunolabeling of fixed embryos or single pulses of calcein at a selection of timepoints (Ettensohn and Malinda, 1993; Guss and Ettensohn, 1997). We found that, while both approaches observed similar dynamics in elongation of skeletal elements (Fig. 3), polychrome labeling was a more direct and accurate method for measuring skeletal growth.

To further explore the temporal information that can be gained from simple polychrome-labeling experiments, we examined fluorochrome distribution in the triradiates of control and pharmacologically or chemically perturbed embryos (Fig. 4; Fig. 5). When we compared fluorochrome distribution in the triradiates of embryos treated with chlorate, axitinib, or SB43 to that in controls, we see that the timing and synchrony of triradiate formation is not affected (Fig. 4C-E, compare to Fig. 2B). This is not surprising, since each of these perturbants is administered at or after 15 hpf, when the triradiates have been initiated (Fig. 5B, K). Furthermore, this observation agrees with what would be expected based on the specific events in skeletal patterning disrupted by each drug. Chlorate treatment at early gastrula stage results in loss of primary ventral elements (Piacentino et al., 2016a) that arise following triradiate formation. It is therefore plausible that triradiate formation is unaffected by late chlorate treatment. Axitinib, a VEGFR inhibitor, produces stunted, abnormally rotated skeletons with missing secondary elements when administered during gastrulation (Adomako-Ankomah and Ettensohn, 2013; Piacentino et al., 2016a)(Movie S2). We show here that axitinib treatment is not sufficient to perturb triradiate timing or synchrony, as demonstrated by clear xylenol orange labeling of both triradiate spicules by 24 hpf (Fig. 4D; Movie S4, compare to Movie S3). This is unsurprisingly, considering that axitinib treatment occurs at 16 hpf, which is just after the triradiates would have initiated (Fig. 5B, K). This is also consistent with previous observations that axitinib treatment does not block the formation of the ventrolateral clusters of PMCs, which secrete the initial triradiates (Adomako-Ankomah & Ettensohn, 2013). Finally, SB43 affects anterior skeletal patterning (Piacentino et al., 2015). Our observation that triradiate formation is unaffected by SB43 (Fig. 4E) is consistent with our expectations, since formation of anterior skeletal elements is a relatively late event that occurs well after the triradiates have formed and elongated.

Of the four perturbations explored in this study, nickel treatment was the only condition that showed dramatic effects on the timing of triradiate formation. In order to understand this effect more clearly, we first established that triradiate formation initiates synchronously in control embryos around 15 hpf (Fig. 5B, K). This was a surprising result, since traditional birefringent imaging methods first detect the spicules of the triradiates after mid-gastrula stage (Decker et al., 1988; Killian and Wilt, 2017). This discrepancy suggests that calcium-binding fluorochromes offer a more sensitive approach for the detection of skeleton biomineral than birefringence, particularly for early events in skeletogenesis.

We next compared the normal temporal dynamics of triradiate initiation to the dynamics in nickel-treated embryos. Our results reveal that NiCl_2_ treatment is sufficient to both delay and desynchronize triradiate formation (Fig. 4B; Fig. 5), which is a novel observation. Interestingly, this was not a nickel-specific effect: embryos ventralized by early chlorate treatment also produced radial triradiates asynchronously, though the delay in triradiate initiation in chlorate-treated embryos was approximately three hours shorter than the delay in nickel-treated embryos (Fig. S3). The desynchronization of triradiate formation suggests that the ectoderm in embryos ventralized by nickel or chlorate treatment is heterogenous, which is unexpected. Since nickel and chlorate treatment each induce broad ventralization of the ectoderm, we would have anticipated that the ectoderm is uniform; however, our observations of extended asynchronous initiation of triradiates suggests that different clusters of PMCs receive the signal to initiate skeletogenesis at different times. Sulfated proteoglycans (sPGs) are normally present in the ventral and ventrolateral ectoderm adjacent to triradiate-producing PMC clusters and are required during gastrulation for ventral PMC migration and ventral skeletogenesis (Piacentino et al., 2016a). Chlorate treatment inhibits sPG formation by blocking the addition of sulfates (Bergeron et al., 2011; Piacentino et al., 2016a). Our findings herein reveal that sPGs are also required for timely and synchronous triradiate formation.

The sequential appearance of spatially adjacent triradiates in ventralized embryos implies that some type of communication occurs between a triradiate-producing PMC cluster and its neighbors to favor proximal initiation of an adjacent triradiate, potentially through the titration of VEGF signals (Ettensohn and Adomako-Ankomah, 2019). Unfortunately, we do not understand the signals that promote triradiate formation: to date, no patterning cues have been identified that specifically block triradiate formation without also blocking overall biomineralization. Thus, further studies will be necessary to determine the mechanism(s) by which the triradiates form sequentially rather than synchronously over a broad time range in ventralized embryos and why there is a temporal difference in the delay of triradiate initiation in nickel- and chlorate-treated embryos.

Although axitinib treatment did not affect the timing or synchrony of triradiate formation (Fig. 4D; Movie S4, compare to Movie S3), VEGFR inhibition resulted in abnormal orientation of the triradiates and reduce elongation rates for several skeletal elements. Previous studies have found that axitinib treatment results in dramatic stunting of the oral and aboral skeletal rods (Adomako-Ankomah and Ettensohn, 2013; Piacentino et al., 2016a). Here, we confirm and quantify the significant stunting of the oral and aboral rods in axitinib-treated embryos compared to controls and show that this defect arises by 27 hpf. We also show that there is significant stunting of the ventral transverse rods as early as 21 hpf, as well as mild but statistically significant stunting of the dorsal-ventral connecting rods, body rods, and recurrent rods by 27 hpf (Fig. 6A-D; Table 1; Table S2; Movie S6, compare to Movie S5). Thus, axitinib treatment was sufficient to dramatically slow, but not completely halt, the elongation of most skeletal elements after 21 hpf. These observations are consistent with the requirement of VEGF signaling for PMC migration and biomineralization.

Additionally, our lab has previously observed that the body rods in axitinib-treated embryos are abnormally oriented relative to the midline (Piacentino et al., 2016a). Here we explore this anterior-posterior orientation defect further and find that this is not due to structural changes in the shape or branching of the triradiates themselves (Fig. 6E-G), but rather due to abnormal orientation of the triradiates relative to the midline (Fig. 6G, H). This is an interesting result since axitinib was added to the larval cultures at 16 hpf (Sun and Ettensohn, 2014), slightly after the initiation of triradiate formation at 15 hpf (Fig. 5B, K). This finding suggests that the orientation of the triradiates is still being determined even after biomineralization has occurred, which agrees with previous reports in *Psammechinus miliaris* that the triradiate rotational positions are not initially fixed and they instead continue to rotate, similar to the rotation of a propeller, during early stages of skeletogenesis (Wolpert and Gustafson, 1961). Furthermore, we found that the left and right triradiates in axitinib-treated embryos were oriented at different degrees of rotation relative to the midline, suggesting a perturbation of bilateral symmetry in these embryos. Our findings suggest that VEGF signaling contributes to symmetrically orienting the triradiate rotational positions relative to the overall body axes.

Here we have established a polychrome labeling approach with calcium-binding fluorochromes in nested pulse-chase experiments to gain unique temporal insight into control and perturbed skeletal patterning in sea urchin embryos. We use that approach to reveal temporal asymmetry of triradiate formation in ventralized embryos, and abnormal triradiate orientation in VEGFR-inhibited embryos. We anticipate expansion of the applications for this approach by our group and others, including three-or four-pulse labeling of embryos with skeletal patterning perturbations to determine the etiology of those defects.

## Materials and methods

### Animals, embryo cultures, imaging

Adult *L. variegatus* sea urchins were obtained from either the Duke University Marine Labs (Beaufort, NC) or Reeftopia (Miami, FL). Gametes were collected into ASW from adults by intraperitoneal injection of 0.5 M potassium chloride. Embryos were cultured at 23°C in artificial sea water (ASW) with the listed concentrations of fluorochromes and drugs, where applicable. Embryos were imaged on a Zeiss Axioplan upright microscope at 200 x magnification with Differential Interference Contrast (DIC) to visualize morphology or at multiple focal planes with plane-polarized light to visualize birefringent skeletons. Skeleton images were montaged in ImageJ and excess out-of-focus light was manually removed so that the entire skeleton was in focus.

### Mineralization marker staining

Alizarin red S (Sigma #A5533), xylenol orange tetrasodium salt (Sigma #398187), calcein (green) (ThermoFisher #C481), and calcein blue (Sigma #M1255) stocks were separately prepared by dissolving each chemical in distilled water. Tetracycline hydrochloride (Sigma #T3383) stock was prepared by dissolving the chemical in 70% ethanol. Unless otherwise noted, embryos were incubated in 30 μM alizarin red (AZ), 30 μM xylenol orange (XO), 60 μM tetracycline (TET), 15 μM calcein green (CG), or 45 μM calcein blue (CB) for the time periods specified. For polychrome labeling, embryos were knocked down with 2X ASW and washed thoroughly with 1X ASW at least three times before the next fluorochrome was added.

### Drug treatments

Embryos were incubated in 0.5 mM nickel chloride (Sigma #339350) from fertilization to early gastrula stage (Hardin et al., 1992), 20 mM sodium chlorate (Sigma #244147) from fertilization to early gastrula stage to ventralize embryos (Bergeron et al., 2011), or from early gastrula stage until imaging to elicit skeletal patterning defects (Piacentino et al., 2016a), 40 nM axitinib (Sigma #PZ0193) from 16 hpf until imaging (Sun and Ettensohn, 2014), or 3 μM SB431542 (Sigma #S4317) from early gastrula stage until imaging (Bradham and McClay, 2006) as previously described.

### Microscopy and z-normalization

Differential interference contrast (DIC) and birefringence (plane-polarized light) images were collected using a Zeiss Axioplan microscope at 200x. Confocal z-stacks were collected using a Nikon C2Si laser-scanning confocal microscope or an Olympus FV10i laser-scanning confocal microscope at 400x. Lasers used were 561 nm for Alizarin red and xylenol orange, 588 nm for Tetracycline and calcein green, or 405 nm for calcein blue. Maximum intensity 2-D or 3-D projections were created using ImageJ or Napari. Where necessary, z-normalization was performed by applying a custom *normalize*.*py* script in Python (https://github.com/BradhamLab/image_scripts).

### Scoring, measurements, statistics

Each experiment was replicated at least three times using biologically independent samples unless otherwise noted; the n for each experiment is indicated. Measurements of rod lengths and the angles between skeletal rods in live embryos were performed by rotating 3-D image projections in ImageJ to optimize inspection of the rod(s) of interest, then measuring the length of the rod or inter-rod angles with built-in tools. Rod elongation rates were determined by measuring extent of fluorochrome incorporation from nested pulse-chase experiments. For fixed embryos, rod length measurements were performed using the spatial coordinates of rod endpoints from confocal z-stacks, then calculating from them. For some comparisons, the midline was defined as a line intersecting the center of the blastopore/gut and the dorsal apex/scheitel. Statistical significance was analyzed using paired student *t*-tests with 2-tailed, heteroscedastic settings in Microsoft Excel; p-values < 0.05 were considered significant.

## Author contributions

A.E.D. and C.A.B conceived the study. A.E.D. executed the fluorochrome labeling experiments, and D.T.Z. provided the measurements from fixed embryos for Figure 3 A-G (orange). A.E.D. and C.A.B. analyzed the results and wrote the manuscript; all authors edited the manuscript.

## Acknowledgements

We thank Todd Blute for confocal training and equipment support and the members of the Bradham lab for helpful discussions and feedback. This work was supported by NSF IOS 1656752 award (CAB); A.E.D. was partially supported by the Kilachand Fellowship from the Boston University Multicellular Design Program.

## Graphical Abstract

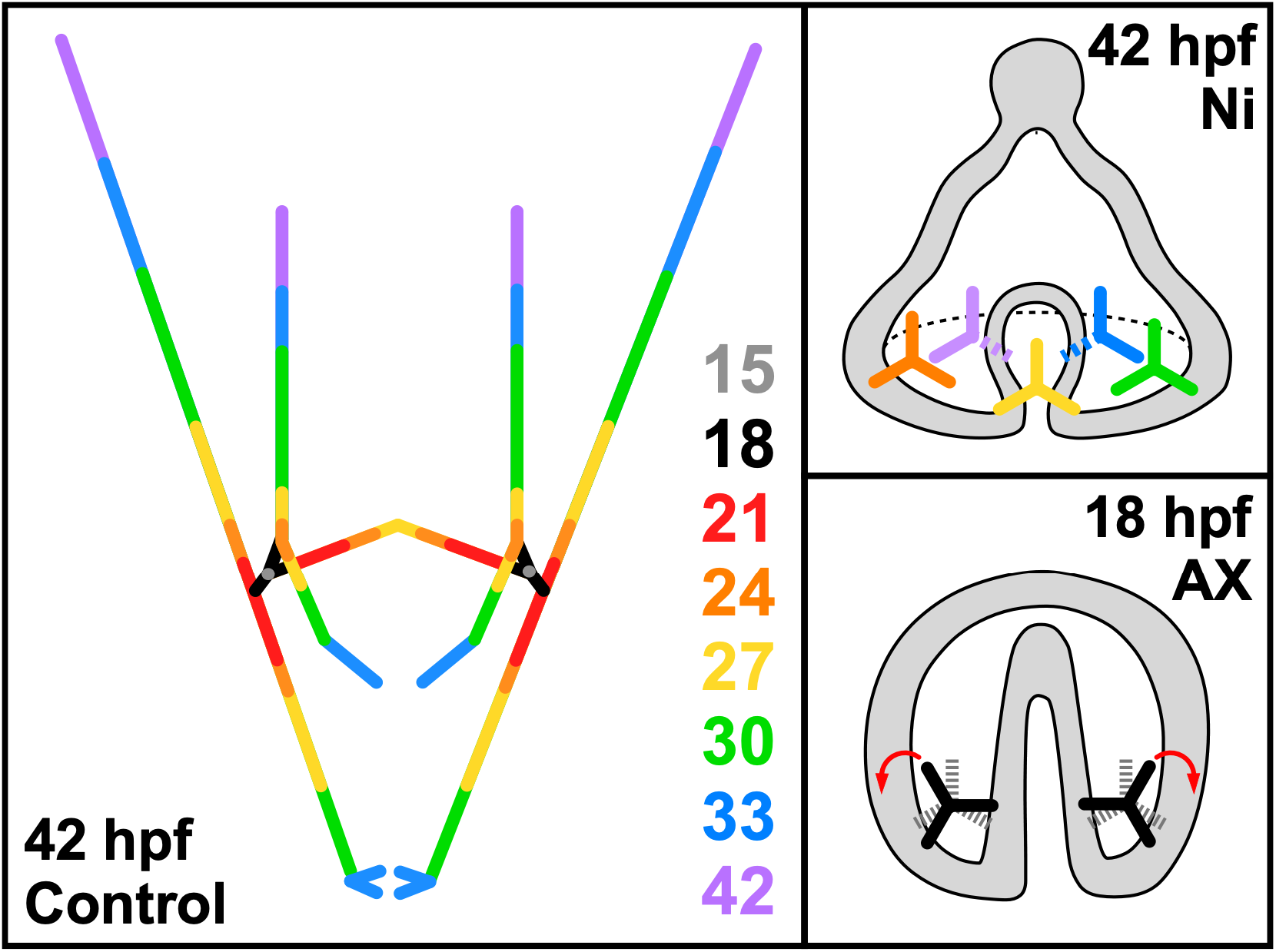

