## Supplemental Tables and Figures for "Polychrome labeling reveals skeletal triradiate and elongation dynamics and abnormalities in patterning cue-perturbed embryos"

**Supplemental Materials for Descoteaux, et al.**

*Contents*

1. Supplemental Tables 1-2

2. Supplemental Figures 1-3

3. Supplemental Movies 1-6

**Supplemental Table 1. Peak excitation and emission wavelengths of three calcium-binding fluorochromes.**

| Fluorochrome | Peak Excitation (nm) | Peak Emission (nm) |
|---|---|---|
| Xylenol Orange | 440 / 570 | 610 |
| Calcein Green | 494 | 517 |
| Calcein Blue | 373 | 420-440 |

**Supplemental Table 2. Skeletal element lengths at three time points in control (C) and axitinib-treated (AX) embryos.**

| | | Anon | VT | DVC | BR | AR | OR | RR |
|---|---|---|---|---|---|---|---|---|
| **C** | **21 hpf** | 14.99 ± 0.24[1] | 45.21 ± 0.70 | 36.90 ± 1.35 | 60.74 ± 1.31 | 25.36 ± 0.79 | 0 | 0 |
| | **27 hpf** | 15.46 ± 0.30 | 46.32 ± 0.85 | 42.18 ± 0.61 | 87.32 ± 1.27 | 59.97 ± 4.98 | 14.86 ± 1.18 | 12.62 ± 0.74 |
| | **42 hpf** | 15.44 ± 0.38 | 46.77 ± 0.99 | 42.62 ± 0.52 | 98.77 ± 1.61 | 218.84 ± 5.83 | 120.12 ± 3.92 | 39.78 ± 3.22 |
| **AX** | **21 hpf** | 15.10 ± 0.22 | 35.04 ± 1.08 | 35.49 ± 1.43 | 57.48 ± 1.94 | 1.17 ± 0.50 | 0.23 ± 0.23 | 1.05 ± 0.53 |
| | **27 hpf** | 15.04 ± 0.23 | 37.41 ± 1.26 | 36.27 ± 1.37 | 71.82 ± 2.81 | 4.45 ± 0.96 | 3.65 ± 0.89 | 6.42 ± 1.22 |
| | **42 hpf** | 15.36 ± 0.21 | 36.85 ± 1.81 | 37.97 ± 1.11 | 83.38 ± 2.93 | 20.76 ± 3.41 | 15.49 ± 2.33 | 25.84 ± 3.08 |

[1]Lengths shown as average µm ± s.e.m.

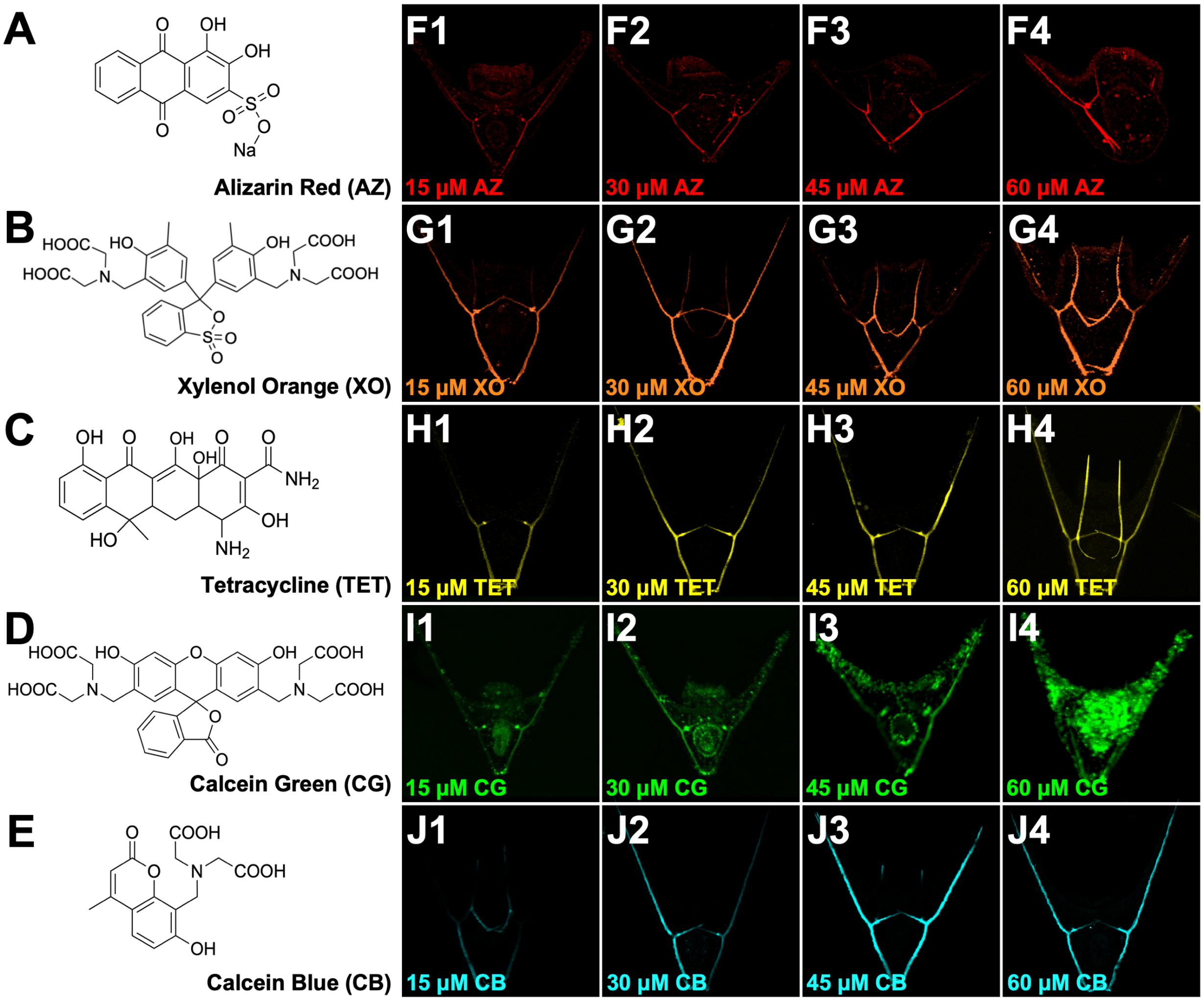


**Supplemental Figure 1. Calcium-binding mineralization markers produce dose-responsive signal levels. A-E.** Chemical structures for five fluorochromes: alizarin red (AZ, A), xylenol orange (XO, B), tetracycline (TET, C), calcein green (CG, D), calcein blue (CB, E). **F-J.** Embryos were incubated in 15 μM (1), 30 μM (2), 45 μM (3), or 60 μM (4) of the indicated fluorochrome from fertilization until confocal imaging at 48 hpf.

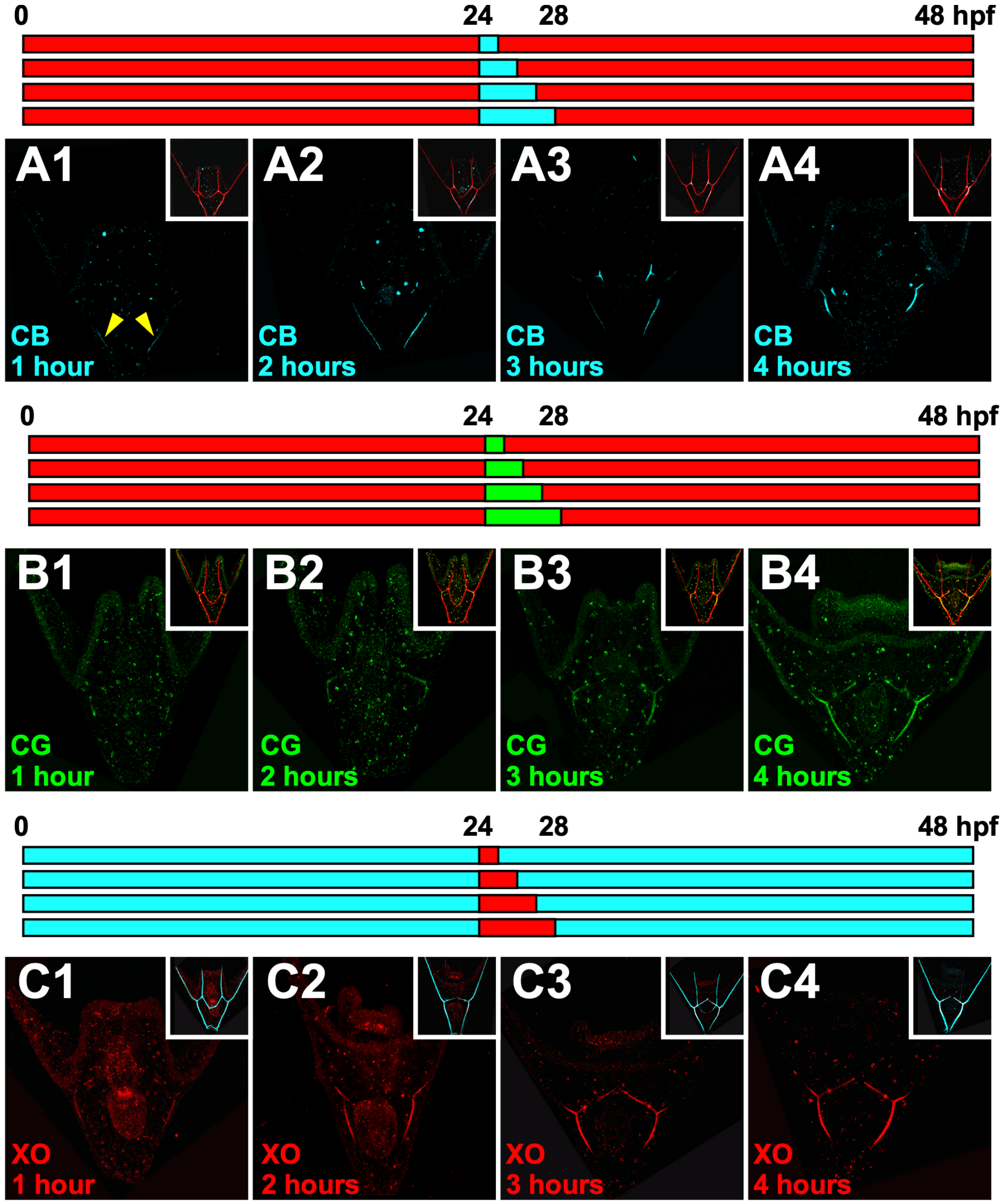


**Supplemental Figure 2. Calcium-binding fluorochromes are most readily detected in the larval skeleton after two to four hours of incubation. A-C.** The schematics (top) illustrate the experimental protocols. Exemplar embryos are shown after labeling with 1- (1), 2- (2), 3- (3), or 4-hour (4) pulses of calcein blue (CB, A), calcein green (CG, B), or xylenol orange (XO, C). Inset shows merged image of the same embryo with XO (A-B) or CB (C) labeling the full skeleton.

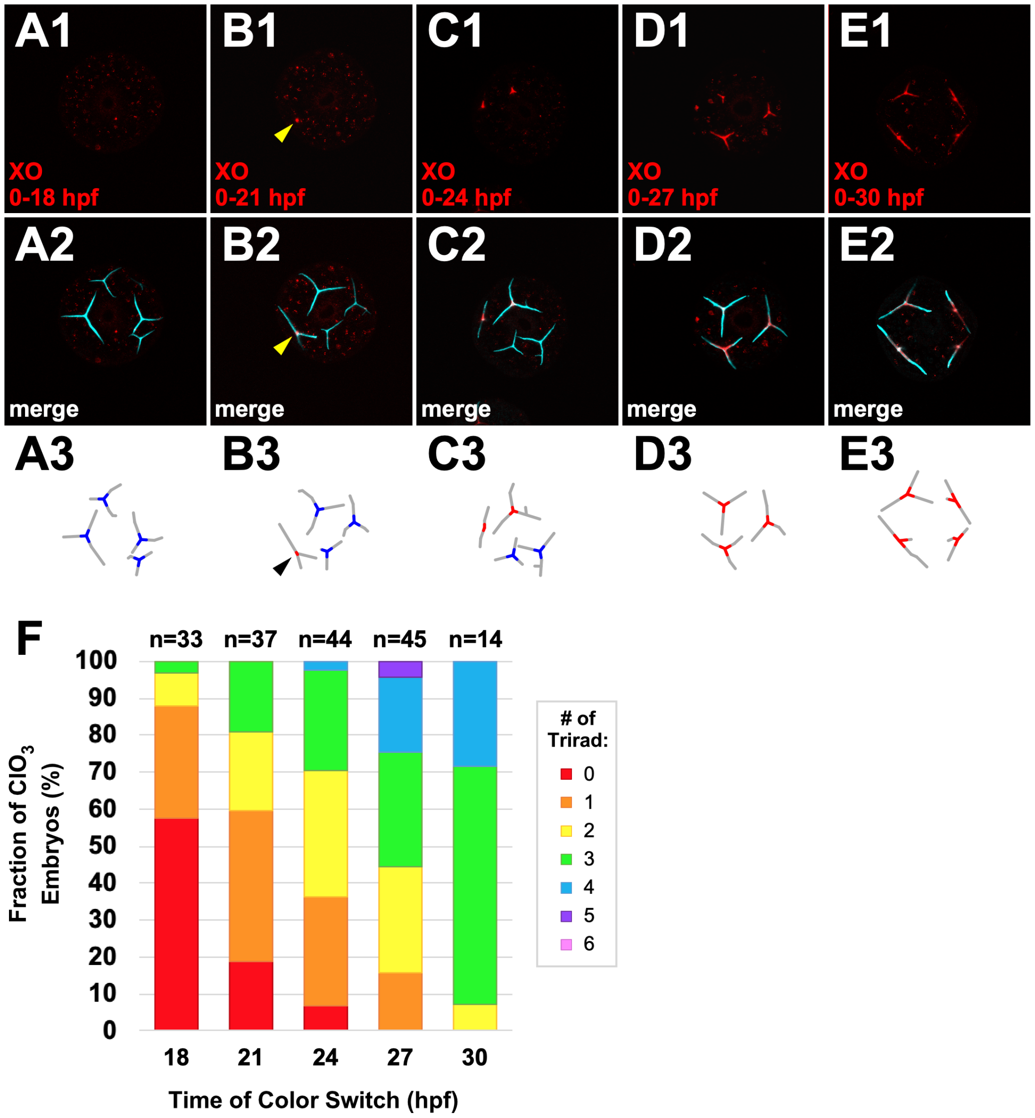


**Supplemental Figure 3. Triradiate formation is delayed and asynchronous in sea urchin embryos radialized by early chlorate treatment. A-E.** Representative confocal images of xylenol orange (1) or merged fluorochrome fluorescence (2) of embryos ventralized with chlorate treatment (20 mM) from 0-15 hpf and dual-labeled with XO and CB that were switched at the indicated timepoints. Schematics (3) show earliest label detected in each triradiate (red = xylenol orange, blue = calcein blue) as well as overall skeleton (gray). **F.** The average percentage of embryos with the indicated number of triradiates formed at each color switch timepoint is shown.

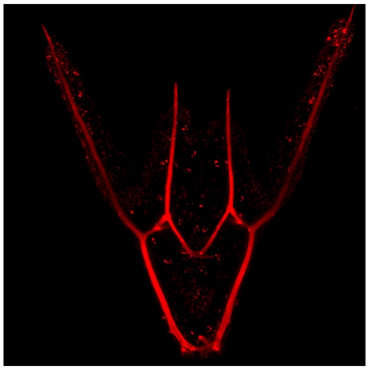

**Movie S1. 3-D rotation of a control embryo labeled with xylenol orange.** 3-D projection of a control embryo labeled with xylenol orange (0-48 hpf) rotating about the dorsal-ventral axis (left) or left-right axis (right). Z-steps are 0.5 μm. Related to Figure 1D.

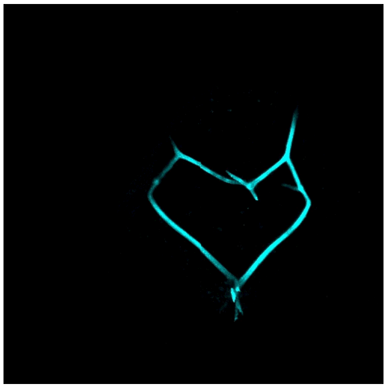

**Movie S2. 3-D rotation of an axitinib embryo labeled with calcein blue.** 3-D projection of an axitinib-treated embryo labeled with xylenol orange (0-48 hpf) rotating about the dorsal-ventral axis (left) or left-right axis (right). Z-steps are 0.5 μm.

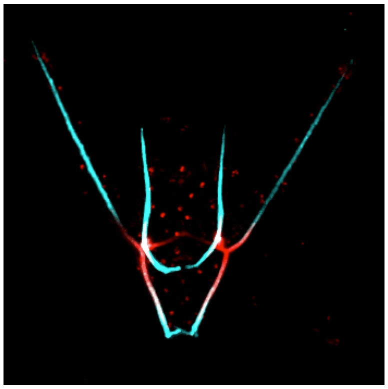

**Movie S3. 3-D rotation of a two-pulse control embryo.** 3-D projection of a control embryo labeled with xylenol orange (0-24 hpf) and calcein blue (24-48 hpf) rotating about the dorsal-ventral axis (left) or left-right axis (right). Z-steps are 0.5 μm. Related to Figure 2B.

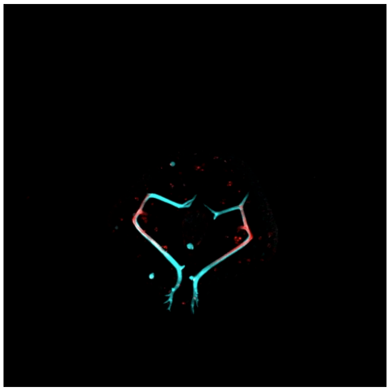

**Movie S4. 3-D rotation of a two-pulse axitinib-treated embryo.** 3-D projection of an axitinib-treated embryo labeled with xylenol orange (0-24 hpf) and calcein blue (24-48 hpf) rotating about the dorsal-ventral axis (left) or left-right axis (right). Z-steps are 0.5 μm. Related to Figure 4D.

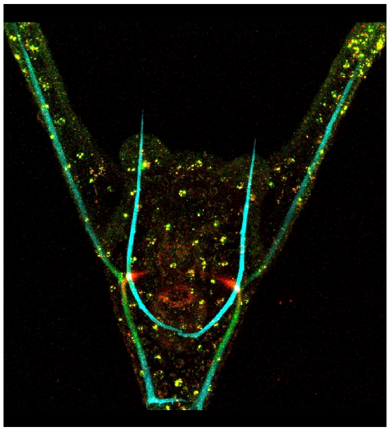

**Movie S5. 3-D rotation of a three-pulse control embryo.** 3-D projection of a control embryo labeled with xylenol orange (0-21 hpf), calcein green (21-27 hpf), and calcein blue (27-42 hpf) rotating about the dorsal-ventral axis (left) or left-right axis (right). Z-steps are 2 μm. Related to Figure 6A.

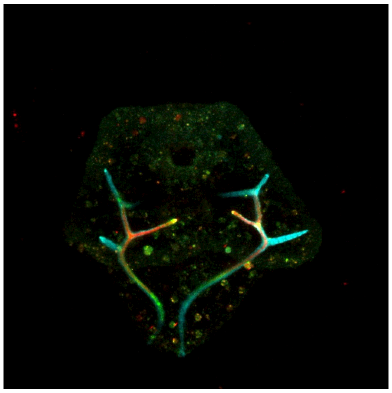

**Movie S6. 3-D rotation of a three-pulse axitinib-treated embryo.** 3-D projection of an axitinib-treated embryo labeled with xylenol orange (0-21 hpf), calcein green (21-27 hpf), and calcein blue (27-42 hpf) rotating about the dorsal-ventral axis (left) or left-right axis (right). Z-steps are 2 μm. Related to Figure 6B.
